# NormiRazor – Tool Applying GPU-accelerated Computing for Determination of Internal References in MicroRNA Transcription Studies

**DOI:** 10.1101/2020.03.11.986901

**Authors:** Szymon Grabia, Ula Smyczynska, Konrad Pagacz, Wojciech Fendler

**Affiliations:** Department of Biostatistics and Translational Medicine, Medical University of Lodz, Lodz 92-215, Poland; Institute of Applied Computer Science, Lodz University of Technology, Lodz 90-537, Poland; Postgraduate School of Molecular Medicine, Medical University of Warsaw, Warsaw 02-091, Poland; Dana-Farber Cancer Institute, Harvard Medical School, Boston, Massachusetts 02115, USA

**Author notes:** Equally contributing (Szymon Grabia, Ula Smyczynska).

## Abstract

**Motivation:** Multi-gene expression assays are an attractive tool in revealing complex regulatory mechanisms in living organisms. Normalization is an indispensable step of data analysis in all those studies, since it removes unwanted, non-biological variability from data. In targeted qPCR assays the normalization is typically performed with respect to prespecified reference genes, but the lack of robust strategy of their selection is reported in literature, especially in studies concerning circulating microRNAs (miRNA).

**Results:** Previous studies concluded that averaged expressions of multi-miRNA combinations are more stable references than single genes. However, due to the number of such combinations the computational load is considerable and may be hindering for objective reference selection in large datasets. Existing implementations of normalization algorithms (geNorm, NormFinder and BestKeeper) have poor performance as every combination is evaluated sequentially. Thus, we designed an integrative tool which implemented those methods in a parallel manner on a graphics processing unit (GPU) using CUDA platform. We tested our approach on publicly available microRNA expression datasets. As a result the times of executions decreased 19-, 105- and 77-fold respectively for geNorm, BestKeeper and NormFinder.

**Availability:** NormiRazor is available as web application at norm.btm.umed.pl.

**Contact:** Wojciech Fendler.

## Introduction

Despite the boom of next-generation sequencing techniques, quantitative RT-PCR (qPCR) assays remain a popular method for evaluating gene expression. Furthermore, due to recent advances in design of highly specific probes this method is becoming effective for analysis of short non-coding RNAs (Jensen *et al.*, 2011). Although such experiments are easy to conduct and non-expensive, they are burdened with a problem of unwanted, non-biological variability in obtained data. Such variability comes mainly from inequality of RNA concentrations, slight differences in samples’ handling and devices’ operation and changes in experimental environment. Since it cannot be avoided at the prelaboratory or laboratory levels, this undesirable component of variation should be removed by a data pre-processing operation, called normalization (Livak and Schmittgen, 2001). Otherwise, those inaccuracies could shadow true biologically relevant differences. Many normalization strategies have been already suggested, but the problem is still considered open to new proposals, especially in those areas where intensive studies have been started only recently (Faraldi *et al.*, 2019; Drobna *et al.*, 2018). One of such fields, that prompted our group to review available normalization algorithms, is research on circulating microRNAs (miRNAs) as potential biomarkers (Fendler *et al.*, 2017; Elias *et al.*, 2017). In past years, circulating miRNAs were shown as very promising diagnostic tools in cancer research (Nakamura et al., 2016; Ogata-Kawata et al., 2014), radiation exposure (Małachowska *et al.*, 2019), diabetes (Satake *et al.*, 2018), cardiology (Nonaka *et al.*, 2019) and numerous other areas. Problems with finding optimal normalization strategy often pose a major obstacle in translating results of research into clinically applicable diagnostic tools (Tiberio *et al.*, 2015; Witwer and Halushka, 2016). In such studies, the first stage usually consists in screening the whole miRNA expression profile in order to identify those miRNAs, which are expressed differentially in the studied condition in comparison to the control group. At this stage, normalization is best performed using a statistical characteristic of samples (e.g. mean expression). It is routinely done when the data comes from high-throughput qPCR (Mohammadian et al., 2013) or RNA sequencing (Zyprych-Walczak et al., 2015).

However, the approach described above is impossible to apply when the qPCR assay targets only few differentially expressed miRNAs (or genes). The mean expression cannot be used as reference in this scenario as the miRNAs are chosen on the basis of between-group differences. Therefore, in such situations, normalization must be performed with respect to endo- or exogenous control genes or total input RNA (Schwarzenbach et al., 2014). Exogenous controls are the RNA (synthetic or coming from other species) that are artificially introduced to the sample in known amounts for the purpose of quality control of consecutive cycles of PCR reaction and normalization procedures. In contrast, endogenous reference genes originate from a tissue sample itself and are selected for their known and experimentally validated, stable expression, in most cases unchanged by pathological conditions or other clinical factors. In miRNA expression experiments either some stably expressed miRNAs or other small RNAs are chosen as those references. However, there is still no consensus on choosing proper internal references for qPCR miRNA studies, especially in biofluids (Marabita *et al.*, 2016; Corral-Vazquez *et al.*, 2017).

Literature review (Drobna et al., 2018; Faraldi et al., 2019; Peltier and Latham, 2008; Sauer et al., 2014) revealed that the most commonly used method of selecting reference miRNAs is the application of algorithms that were previously designed for identification of housekeeping genes for RNA tissue expression studies: geNorm (Vandesompele et al., 2002), NormFinder (Andersen et al., 2004) and BestKeeper (Pfaffl et al., 2002). In these methods potential reference genes are ranked according to their respective stability scores and the most stable one is applied as a normalization factor. Although this approach is commonly used (if any systematic procedure of finding reference genes is applied at all), it has recently been questioned by members of our research team (Pagacz *et al.*, 2020) as an inadequate method of selecting normalization factors.

A new algorithm was therefore proposed that instead of analyzing only single miRNAs, also calculates stability scores for all of their combinations in geNorm (GN), NormFinder (NF or NFG when samples’ groups are provided) and BestKeeper (BK). It was proven that combination-based normalizers are statistically better than single reference genes in terms of stability. Moreover, it was showed that in many cases the most stable reference is formed by averaging expression of 2 or 3 miRNAs that do not necessarily score the highest when analyzed separately. While the gain in finding more stable references is evident, the major obstacle in practical application of this procedure is its computational cost that grows with the number of miRNAs in a dataset and the number of elements in a combination. Initial multi-thread implementation in Python (Pagacz *et al.*, 2020) revealed the problem of long execution times for larger datasets which could not be solved in a fully satisfactory way by CPU parallelization. Following the observation that computations for each of the combinations are independent from one another, we decided to reimplement the proposed algorithm to run on graphical processing units (GPUs). With this idea in mind, we designed a high-speed parallel processing software platform for unbiased combinatorial reference gene selection for normalization of expression data - NormiRazor.

## Implementation

### Implementation for combination-based references

Following (Pagacz *et al.*, 2020), we redesigned the three most popular normalization algorithms in the transcriptomic community, geNorm, BestKeeper and NormFinder, to facilitate calculations of rankings for combination-based references. At the same time, we reimplemented them in Compute Unified Device Architecture (CUDA) model and encapsulated in a self-contained web application called NormiRazor.

The comprehensive combination-based method of identifying reference circulating miRNAs, described in the Introduction section, consists of several operations in the following order:

1. Reading and preprocessing input data.
2. Determination of all possible 2- and 3-element combinations of input miRNAs.
3. Calculation of stability indices for single genes by geNorm, BestKeeper and NormFinder.
4. Calculation of stability indices for combinations by the same algorithms and creating rankings of combinations according to their stability.
5. Aggregation of results from all normalization algorithms and choice of recommended references among combinations.
6. Normalization of input dataset with respect to selected reference (on user request).

Step 1 consists of reading data (gene expression and samples’ group designation), checking format correctness and handling missing values. Step 2 creates the list of all combinations of given length that serves later as a queue of tasks distributed for parallel processing on a GPU. In step 3 single genes are ranked by decreasing stability. Additionally, to reduce computation times for larger datasets (e.g. from NGS), we limit the analysis to 250 genes with the highest stability. Step 4 assesses stability of all combinations and, as the most computationally expensive one, is executed on a GPU. Finally, steps 5-6 summarize the results and allow the user to normalize their data to one of the recommended references. This whole process is also illustrated in Fig. 1A.

**Figure 1:**
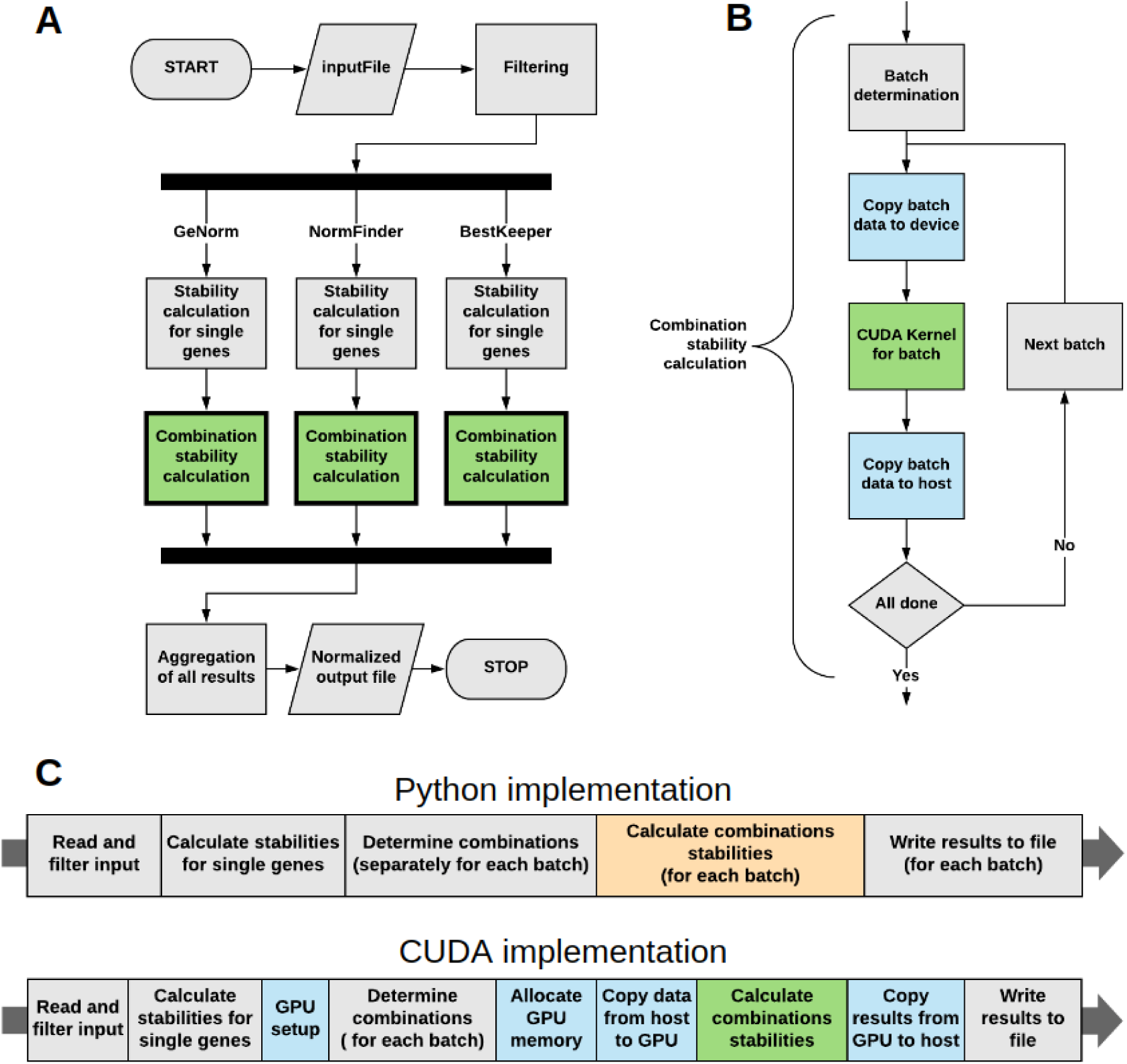
Flow diagram of normalization application execution. A: The whole algorithm; B: A detailed view of operations executed for each batch of input data in CUDA implementation; C: Subdivision of execution time for Python and CUDA implementation for the purpose of benchmarking. Colors: gray – operations on CPU, green – operations on GPU, blue – steps involving both GPU and CPU, orange – operations running on multiple CPU threads.

### Performance optimization

Firstly, the algorithm generates a list of all potential combination-based references. The only optimization for this part with respect to previous Python implementation was rewriting it as a C++ recursive function. Similarly, the aggregation of all previously generated results does not perform very intensive computations, so it is also executed on CPU.

Calculation of stability indices for combinations presented the greatest area of optimization in the form of:

1. Reducing redundant parts of calculations to a single execution (mathematical optimization).
2. Parallel implementation on CUDA-enabled GPU.
3. Limitation of miRNAs set size by the exclusion of those that are the least probable to form stable combinations.

Mathematical optimization consisted mainly in limiting operations repeated in every iteration to only one execution. Formal algebraic description of the algorithms and some minor modifications introduced by us are presented in detail in Appendix. It was also applicable to Python implementation, but it still have not resulted in achievement of reasonably short execution times that researchers looking for reference miRNA would accept. Thus, observing that calculation for each combination are relatively simple, totally independent from one another and identical in terms of computations and differ only by data, we decided to utilize GPU with its SIMD (single instruction, multiple data) processing paradigm. CUDA technology was chosen due to its established position in research applications, especially in bioinformatics and widespread availability of CUDA-enabled GPUs. Finally, we decided to limit number of potential miRNAs to 250, which is sufficient for a typical dataset containing 150-250 miRNA, detectable in majority of samples and as such suitable to serve as reference genes. Still, the dataset of such size generates from 551300 to 2573000 unique 3-miRNA combination to analyze.

We divided the main part of the application into three separate paths, each of them corresponding to the proposed normalization algorithm. We then implemented them as CUDA C kernels embedded into a C++ application (scheme in Fig. 1B), as to achieve maximal performance improvement. In order to further optimize the application, we decided to implement most of the components of the entire procedure on our own with parameters tailored to our needs rather than using more general third-party solutions.

### Results generation in NormiRazor

At the beginning the stability values for single miRNAs are calculated by three normalization algorithms. Since the outputs of those methods are not comparable, for the purpose of selection of the best individual microRNAs, they are normalized to the range [0, 1]. Then, the program calculates their average across the algorithms and on this basis identifies the list of up to 250 most stable candidates that are later used to form the combination-based references.

Combination stability calculation is performed on a GPU and starts with generation of combination-based reference by averaging expression of 2 or 3 miRNAs from the above mentioned list. When stability of each newly formed potential reference is calculated, this combination is treated as an additional gene, appended to the original expression dataset. Resulting stability of a combination and its ranking in the dataset is saved for the comparison with all other combinations of miRNAs. The combinations are sorted by their ranking and then their stability values, in result of which a separate list is generated for each normalization algorithm. New standardized rankings from value range [0,1] are assigned to the combinations based on their position in sorted list, generated by particular algorithm. The results of all algorithms are aggregated by averaging the rankings from all normalization algorithms applied and the list of the best normalizers based on the outcome of this procedure is generated. We decided to present the top 5 combinations to the user and then allow them to normalize their dataset with such a reference, while the complete results are available for download.

Finally, having identified the need for a readily available tool for users with limited experience in programming, we designed graphical user interface (GUI) for our algorithm that is available on our institution’s server at norm.btm.umed.pl.

### Benchmark

We decided that the main measure of efficiency and user-friendliness of our application would simply be total execution time, since it is the most discernible for the end user and ultimately affects their experience the most. As a way of providing an insight into the particular parts of operation of the application, we also measured the execution times of some distinct parts of the algorithm, including purely computational one, named “kernel” in the results section. We performed time of execution tests on datasets either with varying number of miRNAs (test 1) or samples (test 2). In the first set of tests, we used subsets of expression set GSE75389, available in Gene Expression Omnibus (GEO), and we adjusted the number of miRNAs from 50 to 400 with a step of 50, while we always included all 8 available samples. Benchmark with a varying number of samples (10 to 50, step of 10) was based on another GEO dataset - GSE90828. In this second benchmark we included 156 miRNAs without missing data. For both tests, separate experiments were performed for 2- and 3-element references (mean expression of 2 or 3 miRNAs, respectively). Each measurement was repeated 5 times and the result is presented as mean with standard deviation.

For the purpose of testing the performance improvement of the proposed computing method, we employed two benchmarking platforms. The system specifications of both platforms are presented in Suppl. Table 1. We used the Python algorithms implementations from (Pagacz *et al.*, 2020) as a baseline for our comparisons.

Finally, an example of NormiRazor practical application is presented where we attempted a biologically-oriented validation of references found by our application. It is based on the experiment described in (Mestdagh *et al.*, 2009). Dataset GSE121513, which contains miRNA expression profiles obtained by qPCR of 95 neuroblastoma (NB) samples with and without amplification of MYCN gene, was used. MYCN is a transcription factor that binds to miR-17-92 promoter and is known to affect expression of a few miRNAs. According to TransmiR v2.0 (Tong et al., 2019), experimentally validated targets of MYCN are: mir-106a, mir-17, mir-18a, mir-19a, mir-19b, mir-20a and mir-92a. Thus, we can expect that their expression should differ between MYCN-amplified and MYCN-non-amplified samples and observing such difference indicates that correct normalization was applied. We tested this assumption with data normalized to different references, including the ones found by NormiRazor.

All datasets used in tests are publicly available and were generated by qPCR panel assays. Thus, the data were expressed as qPCR quantification cycles (*C*_*q*_) that reflect number of replication cycles after which each miRNA becomes detectable; thus, lower values of *C*_*q*_ indicates higher miRNA expression (Livak and Schmittgen, 2001).

## Results

Comparison of total execution time between CUDA and Python implementation for 3-miRNA references is shown in Fig. 2. We observed that the total execution time for 3-miRNA references and kernel execution time can be accurately modeled by following relations (for both implementations):

- geNorm:

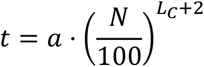
- BestKeeper and NormFinder:

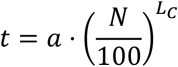

with *N* being the number of miRNAs in a dataset, *L*_*C*_ the number of miRNAs in a combination-based reference and *a* regression coefficient. Division by 100 was included to limit the range of values produced after exponentiation. Curves generated in accordance to the above models are shown in Fig. 2.

**Figure 2:**
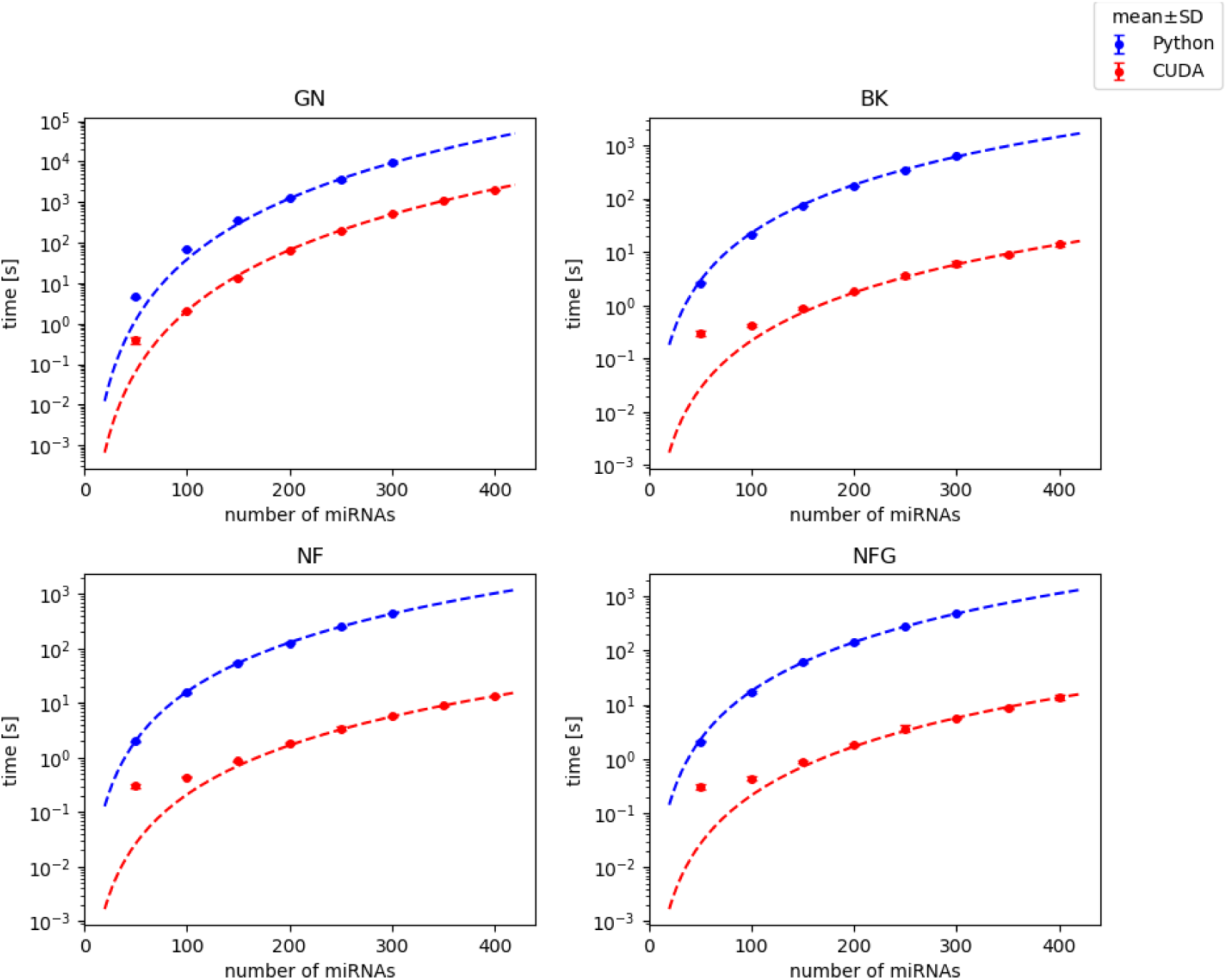
Comparison of total execution time of Python and CUDA implementations for 3-element normalizers on dataset with varying number of miRNAs. Points represent mean and SD of 5 repeated experiments (SD is usually almost negligible), curves are model predictions. Benchmark on platform 1.

For the purpose of quantitative comparison, we defined a speed-up gained by CUDA implementation as:

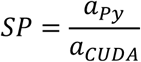

where *a*_*Py*_ and *a*_*CUDA*_ are coefficients from models for Python and CUDA implementations, respectively. Standard deviation (SD) of the speed-up was calculated according to (Dunlap and Silver 1986) as:

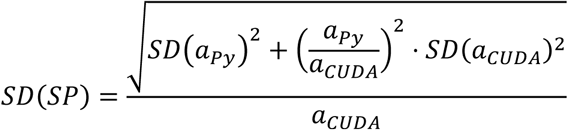

Obtained speed-ups for 3-miRNA combinations were the highest in case of BestKeeper with execution of whole algorithm 100 times faster in CUDA than in Python and the lowest in geNorm with the program still running almost 20 times faster than before (Table 1 and Suppl. Table 2). In the cases of BestKeeper and both versions of NormFinder computational cores (kernels) of algorithms were sped up from about 6000 to almost 22000 times, depending on particular setting, while in the case of geNorm the kernel speed-up was comparable with the total speed-up (Suppl. Fig. 1). A difference of such magnitude was caused by the distribution of execution time between particular operations. From tables in Fig. 3, one can observe that GPU-related operations still take the majority of the execution time of geNorm, which is not the case for BestKeeper and NormFinder. Therefore, the speed-up of the two latter methods affects the whole execution speed-up to a smaller extent than in case of geNorm. The computational core takes the majority of time of GPU operations only in geNorm, while in BestKeeper and NormFinder greater fraction of time is needed for memory allocation and copying data than for actual calculations (Fig. 3). Distribution of execution time between all (CPU and GPU) operations is presented in Suppl. Fig. 2 and for comparison similar plot for Python implementation is shown in Suppl. Fig. 3.

**Table 1:** Speed-up gained by the CUDA implementation with respect to the previous Python version. Benchmark on platform 1.

| Speed-up<br>± SD | total | kernel |  |
| --- | --- | --- | --- |
| Combination length | 3 | 2 | 3 |
| GN | 18.7<br>± 0.6 | 19.3<br>± 1.8 | 18.3<br>± 0.6 |
| BK | 104.7<br>± 4.2 | 12887.5<br>± 1846.3 | 21993.4<br>± 1479.2 |
| NF | 76.5<br>± 2.2 | 6195.2<br>± 1035.1 | 11877.1<br>± 433.5 |
| NFG | 84.0<br>± 3.3 | 6665.7<br>± 1105.0 | 13164.5<br>± 595.6 |

**Figure 3:**
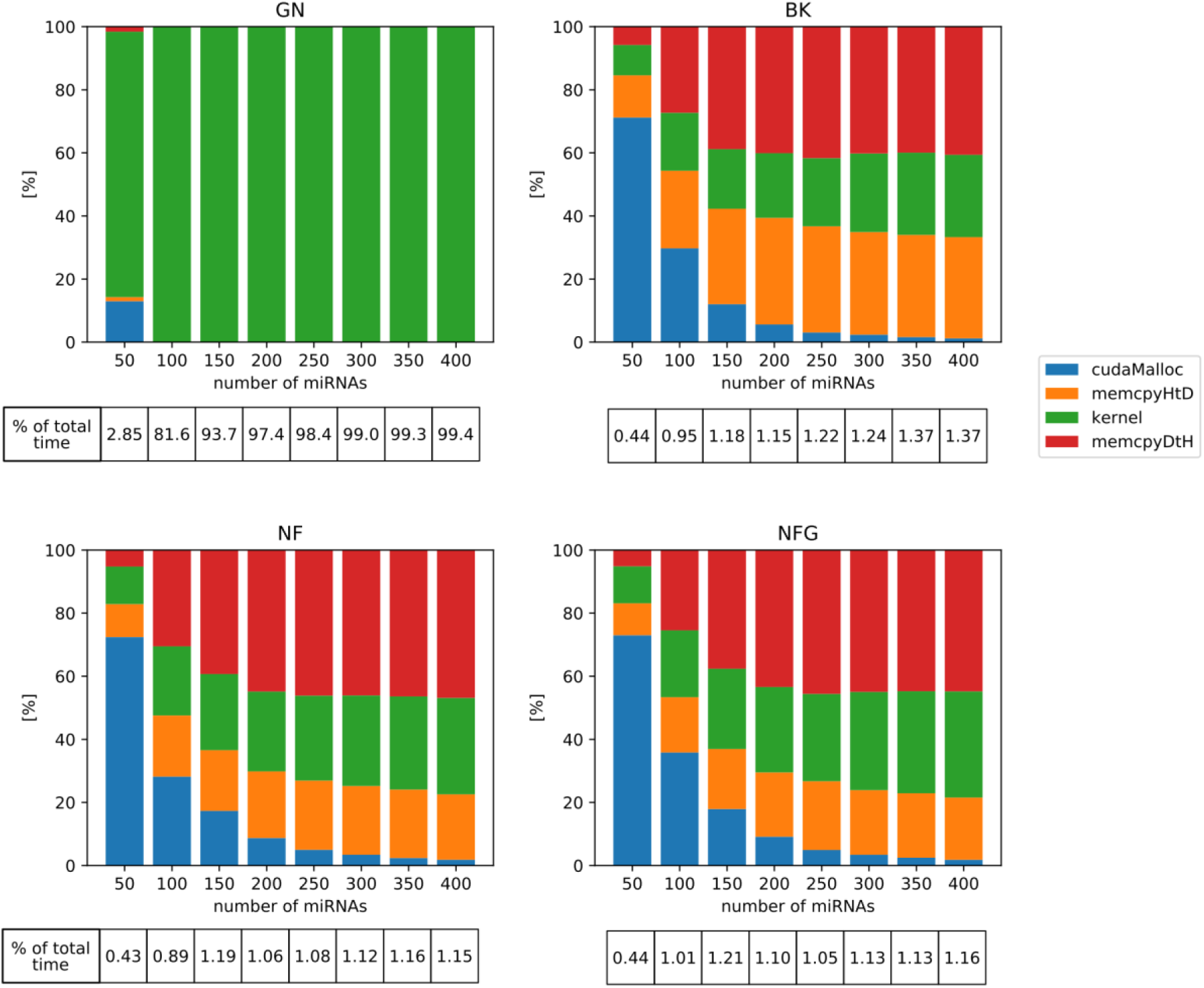
Distribution of execution time between GPU-related operations in CUDA implementations for 3-element normalizers. Tables below the graphs indicate percentage of the total execution time (CPU+GPU) dedicated to operations on GPU. Test 1 on platform 1. cudaMalloc: memory allocation on GPU, memcpyHtD: copying data from RAM to GPU memory, memcpyDtH: copying data from GPU memory to RAM.

Analogical analysis was performed with varying number of samples in dataset (test 2) and the results are shown in Suppl. Fig. 4 and 5. We also performed a comparison of efficiency of GPUs in two benchmark platforms and we obtained very similar kernel execution times in both of them (Suppl. Fig. 6).

Our next goal was to check the validity of identified references when they are applied to real data. Fig. 4 presents the distribution of qPCR *C*_*q*_ values (mean-centered) after different normalization protocols were applied to the data. In panel A, no normalization was applied, thus sample means and range of values were clearly different between samples. The next two panels (B and C) present the effects of normalization to the best 3-miRNA-based reference (averaged expression of miR-1976, miR-30d-5p and miR-214-3p) and normalization to the mean of miRNAs detected in all samples, respectively. Those 2 approaches equalized distributions of *C*_*q*_, such that distributions in all samples became very similar. Additionally, methods B and C do not produce discernibly different results; thus, they may be equally useful. In panels D (original reference 1) and E (original reference 2), references which were proposed in (Schlosser *et al.*, 2015) by the authors of the chosen dataset, were used. Finally, panel F shows detrimental consequences of using a very poor reference (mean of miR-144-3p, miR-1909-3p, miR-451a, identified as the worst by NormiRazor) and highlights how a poor choice in this regard may spawn false positive findings.

**Figure 4:**
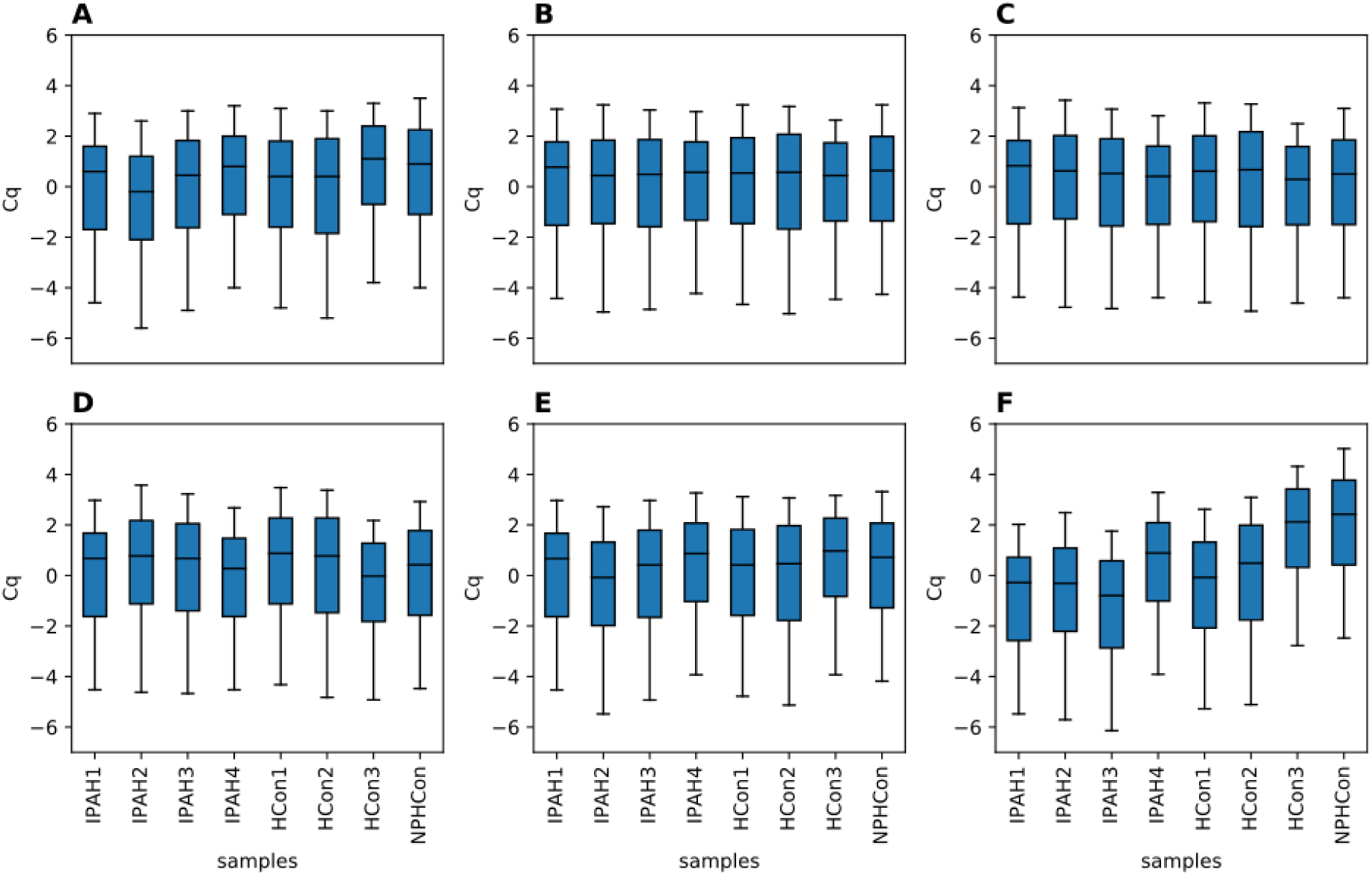
Effect of different normalization schemes on miRNA expression in GSE75389. No normalization (A); normalization: to 3-miRNA reference selected by NormiRazor (B); to mean expression of miRNAs expressed in all samples (C); to references from original article: mean expression of miR-142-3p and miR-320a-3p (D), mean expression of miR-766 and miR-3909 (E); to the worst reference found by NormiRazor (F). Included are only miRNA present in at least 75% of samples; boxes show interquartile range, central line is median, whiskers indicate 5. and 95. percentile.

Effect of normalization methods on variability of miRNA expression between samples was presented in Fig. 5. We observed that normalization to mean expression (gray curve) and normalization to a reference identified by our algorithm (blue curve) produced very similar distributions of expression variances. Moreover, both of them as well as normalization to the original reference 1 (solid green line) were significantly different than lack of normalization plotted in red (p<0.0001 in Kolmogorov-Smirnov test for all comparisons). Standard deviation of expression was on average lowered after of those normalization, excluding purposefully selected worst reference; thus, we conclude that all of these approaches effectively reduce variability. Original reference 2, on the contrary, does not produce distribution different from the one obtained without any normalization (p=0.1201). It is also clear that normalization to poor reference introduces additional amount of variability and is actually worse than lack of normalization (p<0.0001).

**Figure 5:**
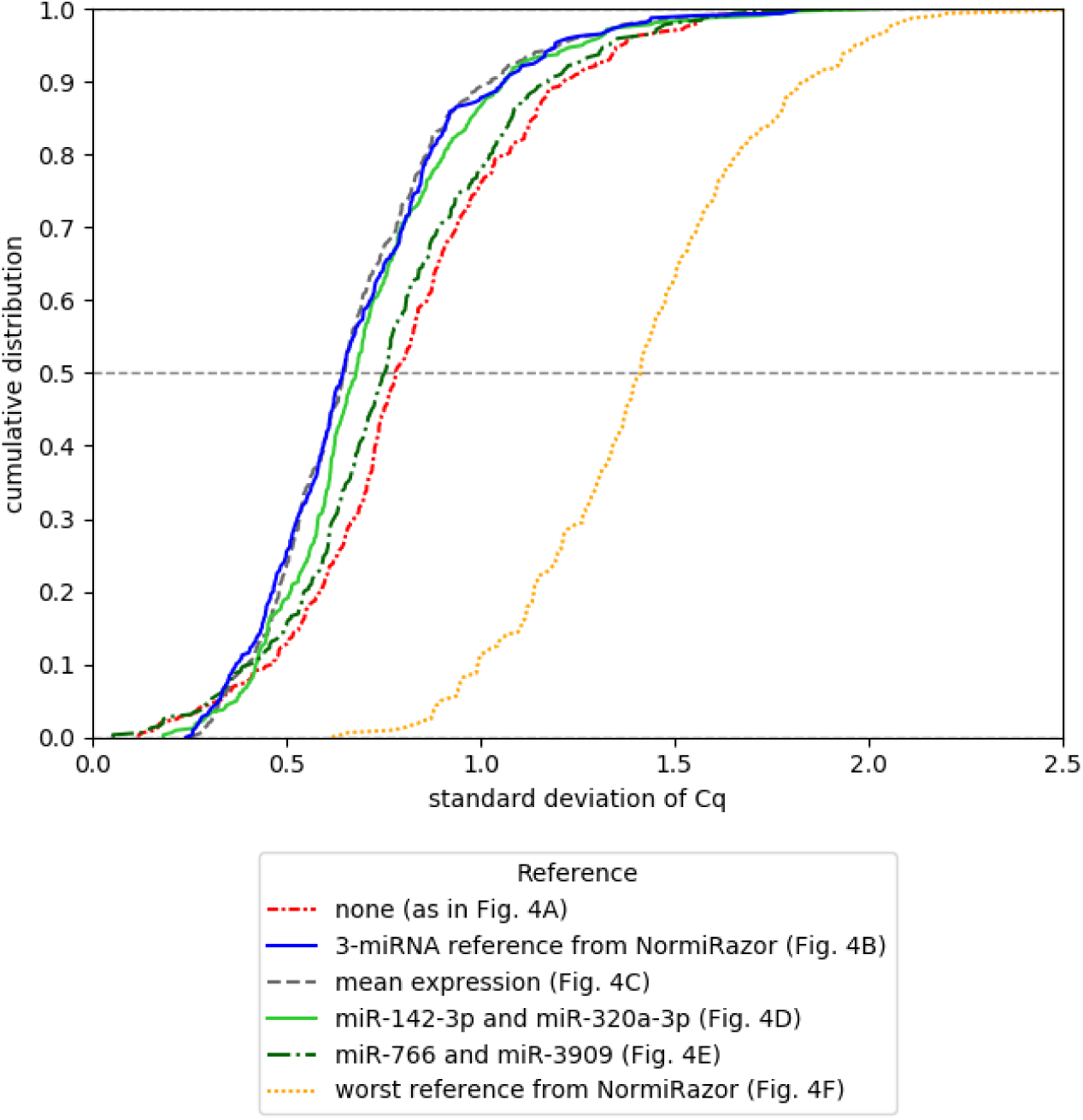
Cumulative distribution of standard deviations of miRNA expression in samples from GSE75389 when different normalization schemes are applied.

Finally, we demonstrated applicability of references found by NormiRazor in differential expression analysis on the exemplary neuroblastoma dataset (GSE121513). The results of this biologically-oriented validation are presented in Fig. 6. We calculated the fold changes (FC) of miRNA expression between MYCN-amplified and non-amplified neuroblastoma samples after applying different normalization schemes. Fig. 6 shows FC values for miRNAs that are known to be affected by MYCN and thus we expected that their expression should be different between two types of neuroblastoma. Whenever statistically significant (FDR-corrected p<0.05) difference was detected, we placed an asterisk above the respective bar. When no normalization was employed, none of miRNAs of interest were found differentially expressed. When a poor reference was tested, only one difference became significant and obtained fold-changes were highly variable. Differential expression analyses after all proper normalizations detected differential expression of 7 out of 9 experimentally validated miRNAs (miR-19a, miR-17-5p, miR-18a, miR-18a*, miR-19b, miR-20a, miR-92). Thus, it must be stated that a proper normalization (either to mean expression or the internal reference) is essential for conclusions of the study.

**Figure 6:**
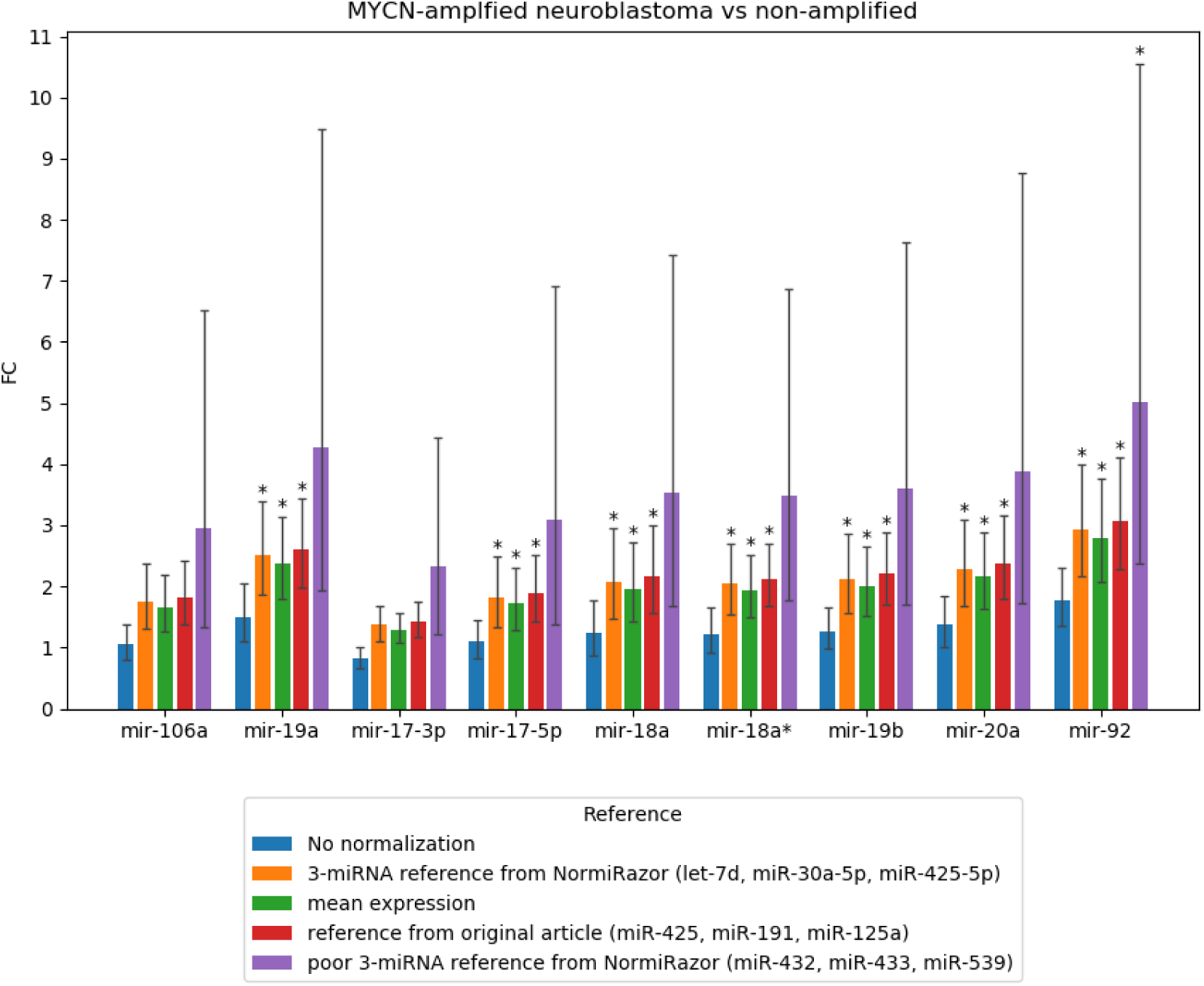
Results of biologically oriented validation on data from GSE121513. Fold-change of miRNA expression between MYCN-amplified and non-amplified neuroblastoma cells is shown with confidence interval when different normalization approaches are applied. Confidence interval is calculated on the assumption that *C*_*q*_ values from qPCR are normally distributed.

## Discussion

In the present study we proposed a new GPU-based computational approach to the recently published (Pagacz *et al.*, 2020) extensive algorithm searching for reference miRNAs. Since the algorithm is based on an analysis of averaged expression of 2 or 3 miRNA as potential reference, it is highly computationally demanding. We identified that majority of calculations are easily parallelizable and can be implemented on a GPU. In our testing environment we observed that the GPU version runs from 20 up to more than 100 times faster than a previous multicore Python implementation. Speed-ups in such a range seem to be typical for GPU reimplementation of bioinformatics algorithms that previously run on CPU. For instance, GBOOST – the tool for a gene-gene interaction analysis – achieved a 40-fold speed-up with respect to its CPU predecessor BOOST (Yung et al., 2011). Even better results were reported by the authors of CUDA-miRanda that is described as 166-times faster than miRanda (Wang et al., 2014). Obviously, interpreting such results, one must take into account characteristics of a particular testing platform (model of CPU and GPU), however, it is still clear that in many applications GPU computing is beneficial.

In the case of searching for reference genes/miRNA, the execution time of an algorithm is an essential part of the user experience. Since finding the reference is only a small, although important part of gene expression experiment, upon which following actions depend, no researcher would accept waiting for the results for an extended period. The longest computational section in our application, namely calculating stabilities for 3-element references by geNorm, takes about 100 seconds with a list of 250 candidate miRNAs, which was set as a technical limit in the final web application. Thus, taking into account the queuing of tasks and aggregation of results, reference miRNAs could be found in about 15 min, which we believe to be an acceptable waiting period. Another important aspect to consider is the quality of identified references. Previous work of our team showed that averaged expression of 2 or 3 miRNA is a more stable reference than any single miRNA with the measure of stability defined as a score from BK, GN and NF (Pagacz *et al.*, 2020). In this study, we additionally showed that references found by NormiRazor effectively reduce variation of miRNA expression. The comparison with the normalization with respect to the mean of all miRNA (considered sometimes as the most reliable one (D’haene et al., 2012)) did not show significant differences in the variability reduction capability.

Many studies stress that normalization strongly affects interpretation of experimental results (Corral-Vazquez et al., 2017; Faraldi et al., 2019; Matoušková et al., 2014). Considering the last stage of our validation, we fully agree with this statement. Differential miRNA expression, that was expected to be observed due to MYCN amplification, was detected when normalization was performed by our algorithm and by the method proposed in the article, upon which this part of validation analysis was based (Mestdagh et al., 2009). When our approach is applied those differences are even more pronounced, so actually a careful, statistical choice of reference miRNA may be even better than the normalization to the mean expression. It seems contradictory to the results reported by Mestdagh et al. (Mestdagh et al., 2009) who argued that the normalization to mean is the most effective method. This may be caused by them comparing their solution only with some limited set of candidate small non-coding RNA (in majority not belonging to the miRNA family), while we scanned all miRNAs expressed in all samples together with their 2- and 3-element combinations.

The entire computing suite is available as a web application. Distributed in this form it is independent of users’ computer hardware and software and it does not require the possession of any CUDA-enabled GPU. Compared with previous solutions, it definitely has several advantages. Initial implementations of geNorm, BestKeeper and NormFinder are already about 15 years old and are not directly compatible with new software. First version of geNorm as VBA applet for Microsoft Excel is no longer available, since according to its website it is “no longer compatible with latest versions of Excel, slow, buggy, difficult to use” (https://genorm.cmgg.be/). Currently geNorm is implemented in commercial qbase+ software, which however is not freely available to the research community. Similarly, BestKeeper cannot be downloaded from its original source. Besides, BestKeeper analysis was originally limited to at most 10 candidate genes that with currently available computational resources is no longer a reasonable limitation. NormFinder macro for Microsoft Excel is the only one that is still available and working even with newest version of Microsoft Office suite. Although it would not be effective for extensive search in combination-based approach, we must acknowledge its stability, that is relatively uncommon in case of bioinformatics tools (Mangul et al., 2019). Some newer implementations have been developed for all the above-mentioned algorithms. NormFinder was rewritten in R and became part of some packages including NormqPCR and SlqPCR. GeNorm and BestKeeper are included in ctrlGene R package. GeNorm belongs also to NormqPCR. They are all in use, but they require R programming skills from their users, who often are biologists or medical doctors rather than data scientists. RefFinder (Xie et al., 2012) is the only freely available tool that combines all 3 algorithms together. A broad review of software packages for identification of reference genes states that all implementations produce consistent results (De Spiegelaere et al., 2015), however they are still capable of analyzing only single genes and not their combinations.

Ultimately, our tool provides researchers working with gene expression data with an easy to use, fast resource for reference variable selection. Its intended and tested field of application are RT-qPCR studies (both panel and targeted) of miRNA expression, however, we are convinced that it can be applied in other omic studies that need to account for technical variability between the samples, similarly to original geNorm, BestKeeper and NormFinder.

## Supporting information

NormiRazor - supplementary results

NormiRazor - appendix

## Acknowledgements

We gratefully acknowledge the support of NVIDIA Corporation with the donation of the Quadro P6000 GPU used for this research.

## References

1. Andersen, C.L. et al. (2004) Normalization of Real-Time quantitative reverse transcription-PCR data: a model-based variance estimation approach to identify genes suited for normalization, applied to bladder and colon cancer data sets. Cancer Res., 64, 5245–5250.

2. Corral-Vazquez, C. et al. (2017) Normalization matters: tracking the best strategy for sperm miRNA quantification. Mol. Hum. Reprod., 23, 45–53.

3. D’haene, B. et al. (2012) miRNA Expression Profiling: From Reference Genes to Global Mean Normalization. In, Next-Generation MicroRNA Expression Profiling Technology: Methods and Protocols, Methods in Molecular Biology., pp. 261–272.

4. Drobna, M. et al. (2018) Identification of Endogenous Control miRNAs for RT-qPCR in T-Cell Acute Lymphoblastic Leukemia. Int. J. Mol. Sci., 19, 2858.

5. Elias, K.M. et al. (2017) Diagnostic potential for a serum miRNA neural network for detection of ovarian cancer. Elife, 6, 1–28.

6. Faraldi, M. et al. (2019) Normalization strategies differently affect circulating miRNA profile associated with the training status. Sci. Rep., 9, 1–13.

7. Fendler, W. et al. (2017) Evolutionarily conserved serum microRNAs predict radiation-induced fatality in nonhuman primates. Sci. Transl. Med., 9, 1–12.

8. Jensen, S.G. et al. (2011) Evaluation of two commercial global miRNA expression profiling platforms for detection of less abundant miRNAs. BMC Genomics, 12, 435.

9. Livak, K.J. and Schmittgen, T.D. (2001) Analysis of Relative Gene Expression Data Using Real-Time Quantitative PCR and the 2-ΔΔCT Method. Methods, 25, 402–408.

10. Małachowska, B. et al. (2019) Circulating microRNAs as Biomarkers of Radiation Exposure: a systematic review and meta-analysis. Int. J. Radiat. Oncol. Biol. Phys., 106, 390–402.

11. Mangul, S. et al. (2019) Challenges and recommendations to improve the installability and archival stability of omics computational tools. PLoS Biol., 17, 1–16.

12. Marabita, F. et al. (2016) Normalization of circulating microRNA expression data obtained by quantitative real-time RT-PCR. Brief. Bioinform., 17, 204–212.

13. Matoušková, P. et al. (2014) Reference genes for real-time PCR quantification of messenger RNAs and microRNAs in mouse model of obesity. PLoS One, 9, 1–11.

14. Mestdagh, P. et al. (2009) A novel and universal method for microRNA RT-qPCR data normalization. Genome Biol., 10, R64.

15. Mohammadian, A. et al. (2013) Normalization of miRNA qPCR high-throughput data: A comparison of methods. Biotechnol. Lett., 35, 843–851.

16. Nakamura, K. et al. (2016) Clinical relevance of circulating cell-free microRNAs in ovarian cancer. Mol. Cancer, 15.

17. Nonaka, C.K.V. et al. (2019) Circulating miRNAs as potential biomarkers associated with cardiac remodeling and fibrosis in chagas disease cardiomyopathy. Int. J. Mol. Sci., 20, 1–16.

18. Ogata-Kawata, H. et al. (2014) Circulating Exosomal microRNAs as Biomarkers of Colon Cancer. PLoS One, 9, e92921.

19. Pagacz, K. et al. (2020) A systemic approach to screening high-throughput RT-qPCR data for a suitable set of reference circulating miRNAs. BMC Genomics, 21, 111.

20. Peltier, H.J. and Latham, G.J. (2008) Normalization of microRNA expression levels in quantitative RT-PCR assays: Identification of suitable reference RNA targets in normal and cancerous human solid tissues. RNA, 14, 844–852.

21. Pfaffl, M.W. et al. (2002) Relative expression software tool (REST) for group-wise comparison and statistical analysis of relative expression results in real-time PCR. Nucleic Acids Res., 30, e36.

22. Satake, E. et al. (2018) Circulating miRNA profiles associated with hyperglycemia in patients with type 1 diabetes. Diabetes, 67, 1013–1023.

23. Sauer, E. et al. (2014) An evidence based strategy for normalization of quantitative PCR data from miRNA expression analysis in forensically relevant body fluids. Forensic Sci. Int. Genet., 11, 174–181.

24. Schlosser, K. et al. (2015) Customized Internal Reference Controls for Improved Assessment of Circulating MicroRNAs in Disease. PLoS One, 10, 1–22.

25. Schwarzenbach, H. et al. (2014) Clinical relevance of circulating cell-free microRNAs in cancer. Nat. Rev. Clin. Oncol., 11, 145–156.

26. De Spiegelaere, W. et al. (2015) Reference gene validation for RT-qPCR, a note on different available software packages. PLoS One, 10, 1–13.

27. Tiberio, P. et al. (2015) Challenges in Using Circulating miRNAs as Cancer Biomarkers. Biomed Res. Int., 2015, 731479.

28. Tong, Z. et al. (2019) TransmiR v2.0: An updated transcription factor-microRNA regulation database. Nucleic Acids Res., 47, D253–D258.

29. Vandesompele, J. et al. (2002) Accurate normalization of real-time quantitative RT-PCR data by geometric averaging of multiple internal control genes. Genome Biol., 3, 0034.1–0034.11.

30. Wang, S. et al. (2014) GAMUT: GPU accelerated microRNA analysis to uncover target genes through CUDA-miRanda. BMC Med. Genomics, 7, S9.

31. Witwer, K.W. and Halushka, M.K. (2016) Toward the promise of microRNAs - Enhancing reproducibility and rigor in microRNA research. RNA Biol., 13, 1103–1116.

32. Xie, F. et al. (2012) miRDeepFinder: A miRNA analysis tool for deep sequencing of plant small RNAs. Plant Mol. Biol., 80, 75–84.

33. Yung, L.S. et al. (2011) GBOOST: A GPU-based tool for detecting gene-gene interactions in genome-wide case control studies. Bioinformatics, 27, 1309–1310.

34. Zyprych-Walczak, J. et al. (2015) The Impact of Normalization Methods on RNA-Seq Data Analysis. Biomed Res. Int., 2015, 621690.

