## Supplementary material for "NormiRazor – Tool Applying GPU-accelerated Computing for Determination of Internal References in MicroRNA Transcription Studies": NormiRazor - supplementary results

**Suppl. Table 1:** Hardware and software configuration of benchmark platforms.

| | **Platform 1** | **Platform 2** |
|---|---|---|
| **CPU** | 2x Intel Xeon E5-2620 v4<br>(8 cores, 16 threads per CPU)<br>2.1 GHz | Intel Xeon E5-2695 v3<br>(14 cores, 28 threads)<br>2.3 GHz |
| **RAM** | 128 GB | 32 GB |
| **GPU** | Nvidia Quadro P6000<br>24 GB GDDR5X<br>3840 CUDA cores | ASUS GeForce GTX 1080Ti<br>11 GB GDDR5X<br>3584 CUDA cores |
| **CUDA Toolkit Version** | 9.1 | 8.0.62 |
| **Python Version** | 3.6.8 | 3.5.2 |
| **Relevant Python modules** | Numpy 1.15.2<br>Scipy 1.0.1<br>Pandas 0.23.4 | Numpy 1.14.5<br>Scipy 1.2.1<br>Pandas 0.24.1 |
| **Threads assigned to Python** | 24 | 16 |

### Additional results of benchmark test 1

**Suppl. Table 2:** Speed-up gained by CUDA implementation with respect to previous Python version. Result from both benchmark platforms.

| Speed-up ± SD | total | | kernel | | | |
|---|---|---|---|---|---|---|
| Comb. len. | 3 | | 2 | | 3 | |
| Platform | 1 | 2 | 1 | 2 | 1 | 2 |
| GN | 18.7 ± 0.6 | 25.6 ± 0.9 | 19.3 ± 1.8 | 26.7 ± 2.2 | 18.3 ± 0.6 | 25.0 ± 0.8 |
| BK | 104.7 ± 4.2 | 153.7 ± 42.1 | 12887.5 ± 1846.3 | 17123.8 ± 1841.3 | 21993.4 ± 1479.2 | 29775.5 ± 1773.2 |
| NF | 76.5 ± 2.2 | 115.0 ± 11.3 | 6195.2 ± 1035.1 | 8798.6 ± 1284.5 | 11877.1 ± 433.5 | 15116.2 ± 564.5 |
| NFG | 84.0 ± 3.3 | 113.9 ± 17.9 | 6665.7 ± 1105.0 | 7887.2 ± 1251.4 | 13164.5 ± 595.6 | 15260.3 ± 558.8 |

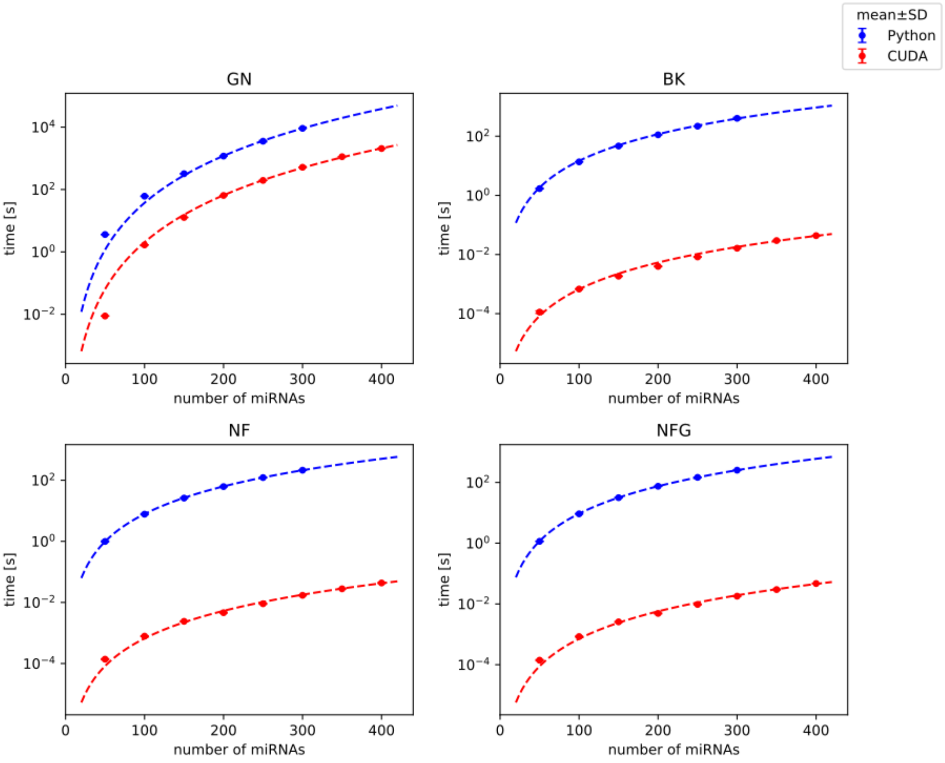


**Suppl. Fig. 1:** Comparison of the calculation (kernel) time of Python and CUDA implementations for 3-element normalizers on datasets with varying number of miRNAs. Benchmark done on the platform 1.

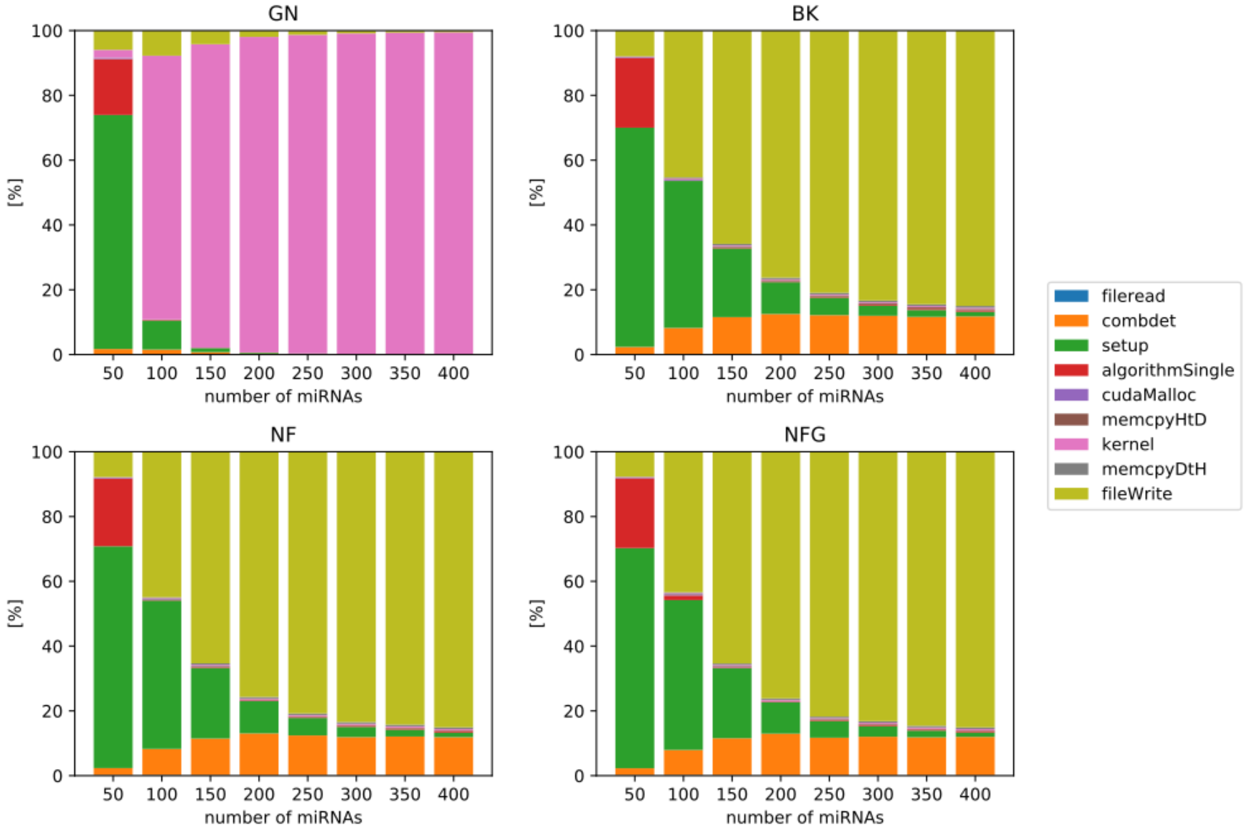


**Suppl. Fig. 2:** Distribution of execution time in CUDA implementations for 3-element normalizers. Test 1 on the platform 1. cudaMalloc: memory allocation on GPU, memcpyHtD – coping data from RAM to GPU memory, memcpyDtH – coping data from GPU memory to RAM, combdet – generation of a combination list, algorithSingle – execution of a given algorithm for single miRNAs.

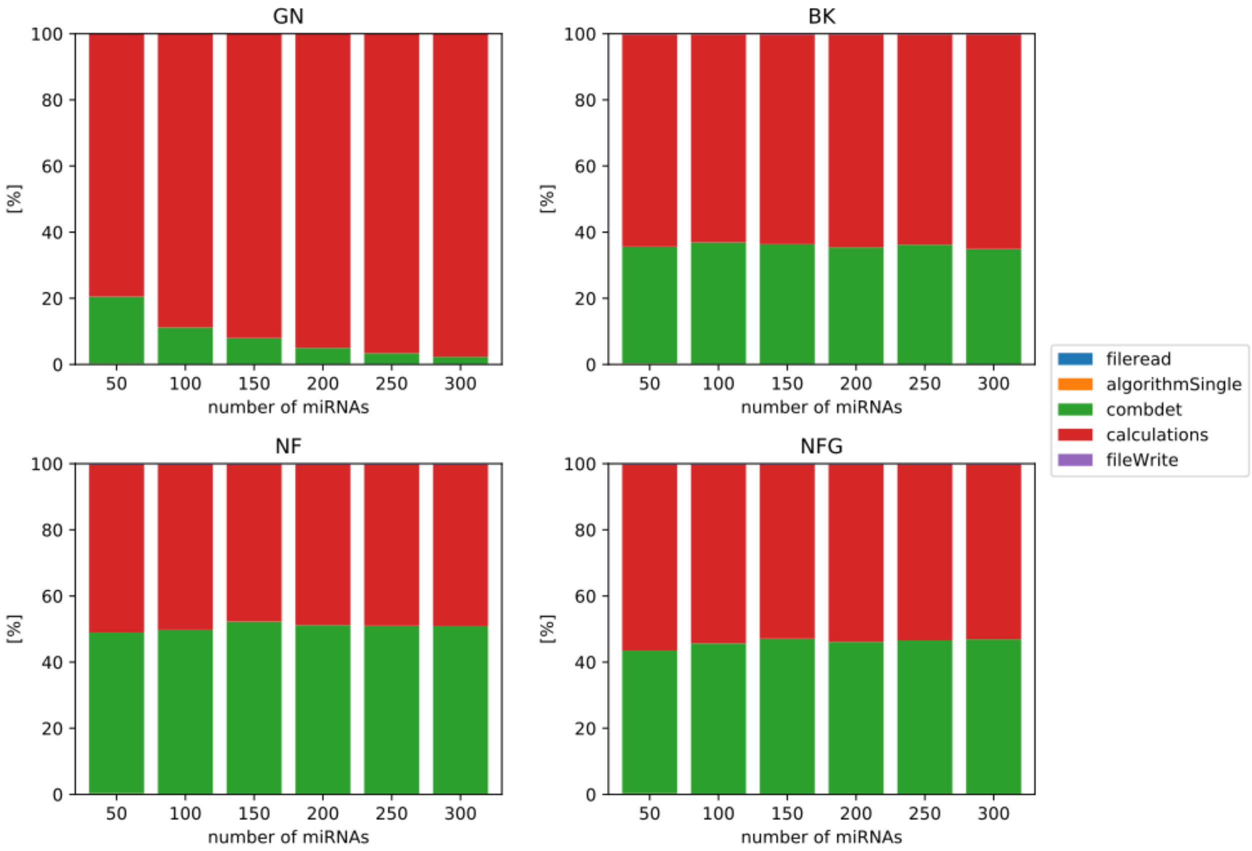


**Suppl. Fig. 3:** Distribution of execution time in Python implementations for 3-element normalizers. Test 1 on the platform 1.

### Results of the benchmark test 2

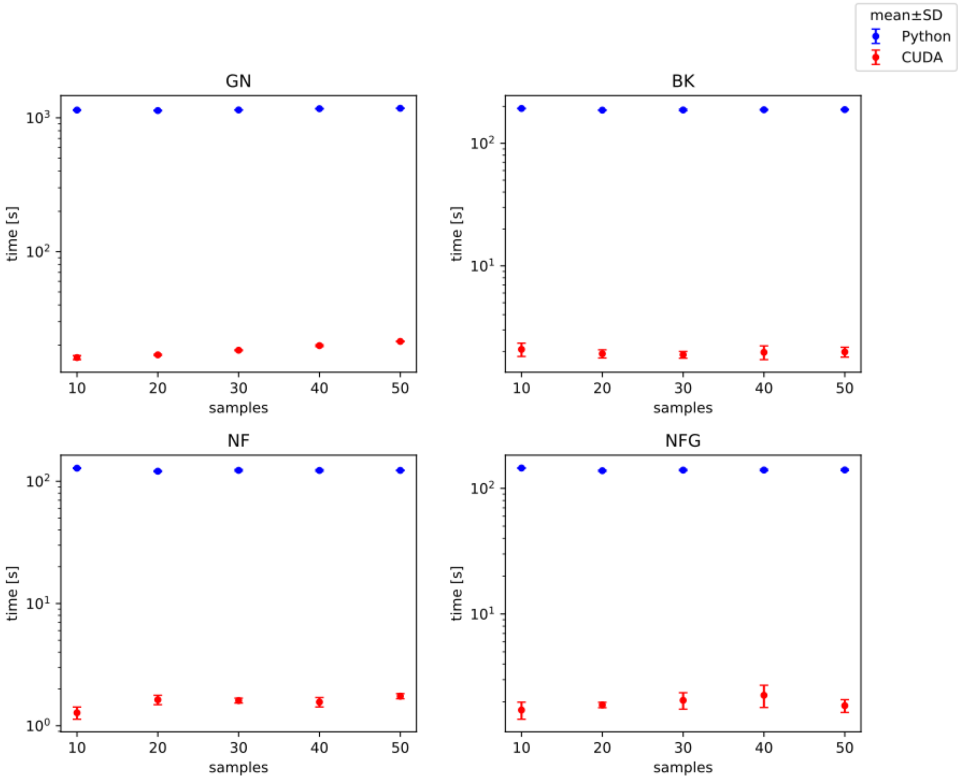


**Suppl. Fig. 4:** Comparison of total execution time of Python and CUDA implementations for 3-element normalizers on dataset with varying number of samples. Benchmark done on the platform 1.

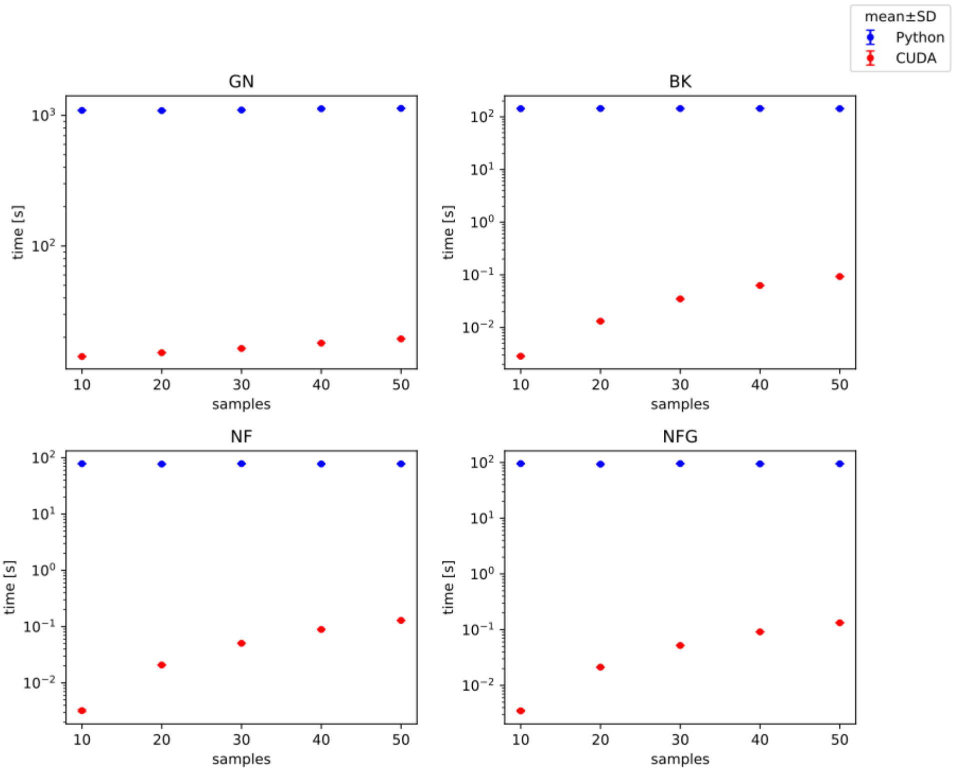


**Suppl. Fig. 5:** Comparison of calculation time of Python and CUDA implementations for 3-element normalizers on dataset with varying number of samples. Benchmark done on the platform 1.

### Comparison of two benchmark platforms

We compared kernel execution time on 2 platforms and plotted the results in Fig. 4. Even though the GPUs that the machines are equipped with are dedicated for different segments of the market, their performance in our case was comparable. Slight advantage of Platform 2 in BK, NF/NFG can be observed, while the Platform 1 was slightly faster with GN.

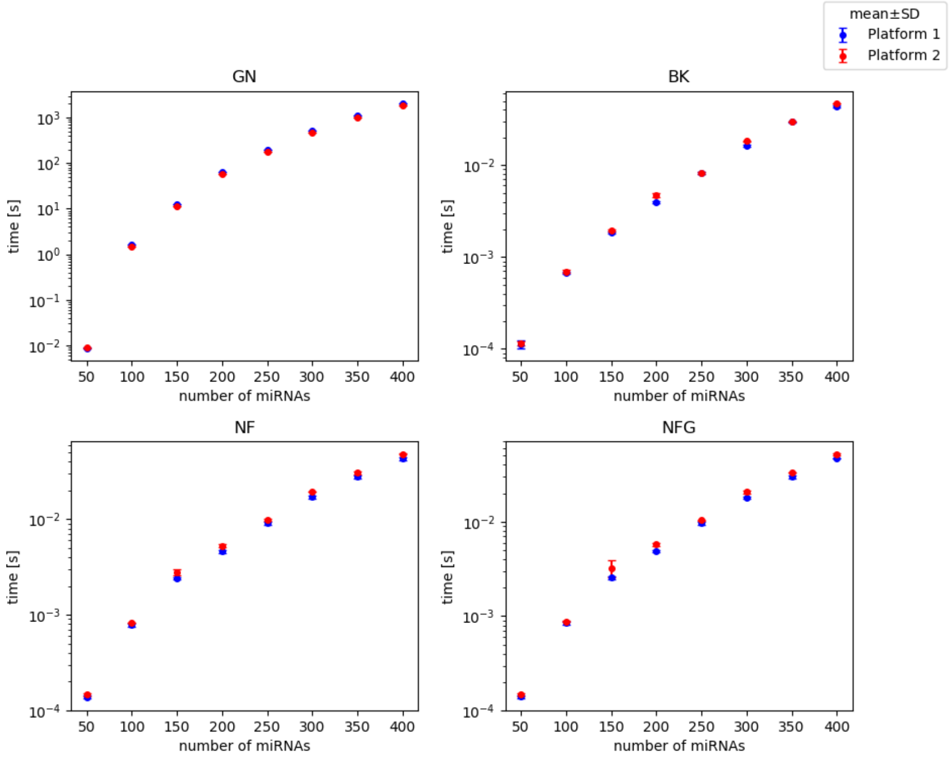


**Suppl. Fig. 6:** Comparison of total execution time on 2 platforms.
