## Supplementary material for "NormiRazor – Tool Applying GPU-accelerated Computing for Determination of Internal References in MicroRNA Transcription Studies": NormiRazor - appendix

In this supplementary file we describe applied normalization algorithms mathematically with modifications and optimization we introduced in order to adjust them to combination-based approach. Designing NormiRazor we decided to standardize input and output of all the algorithms, as well as their mathematical formulation. The following descriptions of normalization algorithms rely on a common data input format, being the gene expression arrays presented below.

Gene expression array (not log-transformed):

$$D = \begin{bmatrix} d_{11} & \dots & d_{1S} \\ \vdots & \ddots & \vdots \\ d_{N1} & \dots & d_{NS} \end{bmatrix}_{NxS} \tag{1}$$

Gene expression array (log-transformed):

$$D_{log} = \begin{bmatrix} log(d_{11}) & \dots & log(d_{1S}) \\ \vdots & \ddots & \vdots \\ log(d_{N1}) & \dots & log(d_{NS}) \end{bmatrix}_{NxS} \tag{2}$$

Array of quantification cycles ($C_q$ values) from qPCR:

$$D_{qPCR} = \begin{bmatrix} C_{q11} & \dots & C_{q1S} \\ \vdots & \ddots & \vdots \\ C_{qN1} & \dots & C_{qNS} \end{bmatrix}_{NxS} \tag{3}$$

where $N$ - number of genes/miRNAs in dataset, $S$ - number of samples.

NormiRazor accepts all of these data formats on the following conditions:

- Correct option is selected at data input.
- Expression array contains only positive values (rows with negative entries are removed from further analysis).

### 1 geNorm

geNorm introduces the internal control gene-stability measure $M$, which represents the stability of any given gene. It assumes that if the ratio of expression values of two genes is constant across all samples, then they can be considered stable and thus used as optimal internal references. Likewise, noticeable variation of the expression ratios indicates the lack of expression constancy in one or both candidate genes.

The algorithm is based on the iterative strategy, where in each iteration the worst candidate for the reference gene is excluded from the dataset. The loop ends when only two genes remain. The final ranking is formed based on the order of genes removal.

During each iteration, first, a 3D matrix of expression ratios of all candidates is determined as:

$$A[n,k,s] = log_2\left(\frac{D[n,s]}{D[k,s]}\right) \tag{4}$$

In case of expression data obtained from qPCR, the input values are already log-scaled and therefore the ratio matrix can be simply calculated as the difference of those values:

$$A[n,k,s] = -(D_{qPCR}[n,s] - D_{qPCR}[k,s]) \tag{5}$$

In this formula, the ratio values are inverted due to the specifics of qPCR method output, in which higher value of quantification cycle indicates lower gene expression.

Then the matrix of pairwise variation $V$, is calculated over all of the samples used in the experiment:

$$V[n,k] = \sqrt{\frac{\sum_{s=1}^{S}(A[n,k,s] - \overline{A}[n,k])^2}{S-1}} \tag{6}$$

where:

$$\overline{A}[n,k] = \frac{\sum_{s=1}^{S} A[n,k,s]}{S} \tag{7}$$

Finally, the $M$-value is calculated for each gene in the dataset:

$$M(i)[n] = \frac{\sum_{k=1}^{N(i)} V[n,k]}{N} \tag{8}$$

where $i$ is the iteration number and $N(i)$ is the number of genes remaining in the dataset during $i^{th}$ iteration of the algorithm.

The gene with the highest value of $M$ is removed from the set and its stability is calculated as the average of all $M$ values before the removal:

$$SI[i] = \frac{\sum_{n=1}^{N(i)} M[n]}{N(i)} \tag{9}$$

Since the values in the matrices $A$ and $V$ remain the same throughout the loop, their determination in each iteration proved to be counterproductive. Therefore, both of them are calculated only once, before the start of the removal procedure. Then in each iteration the next value of $M$ is computed with use of those matrices and the $M$-value from the previous iteration.

$$M(i)[n] = \frac{M(i-1)[n] \cdot N(i-1) - V[n,x]}{N(i)} \tag{10}$$

where $x$ is the index of the removed gene.

All of the steps are summarized in figure below:

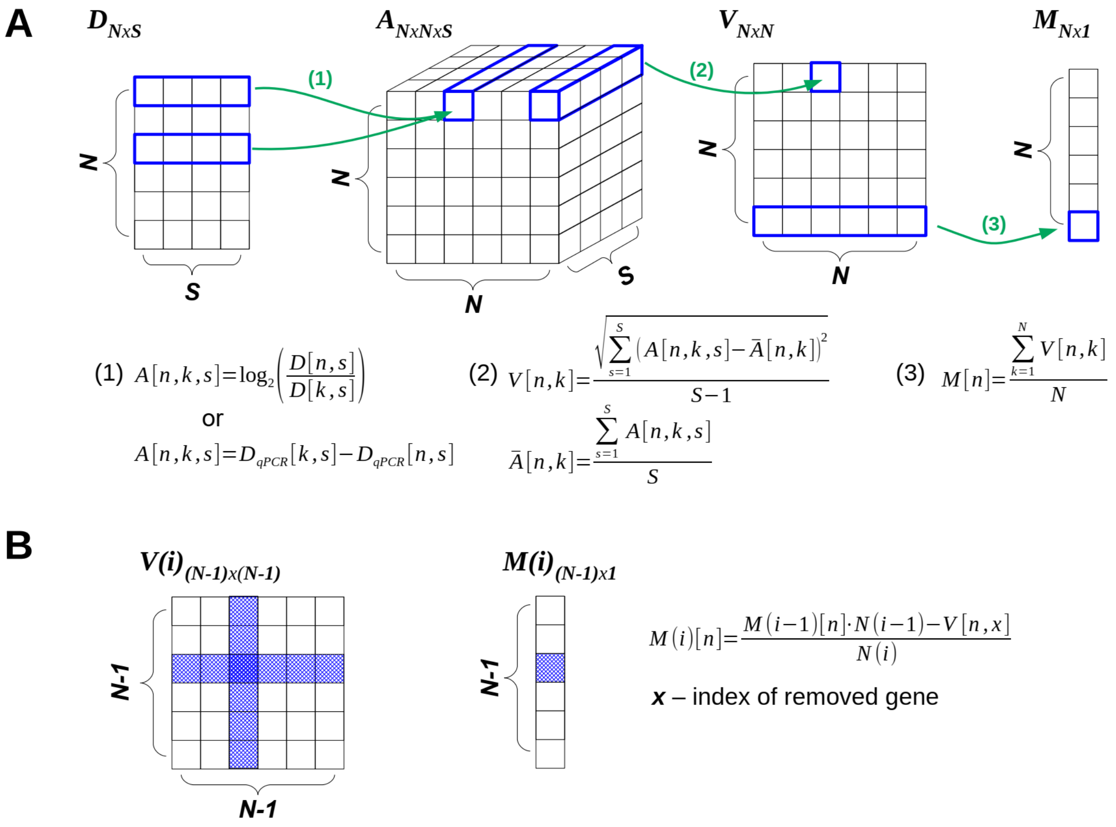

#### 1.1 BestKeeper

The next normalization algorithm, namely BestKeeper, was proposed for the first time as a method for determination of Housekeeping Genes (HKG) with the most stable expressions. Those genes could in turn be combined into an index and used as the baseline for correlations with Target Genes (TG) during an experiment. According to the proposed approach, the geometric mean of all stably expressed genes (called Best Keeper Index ($BKI$)) may act as a good reference in the given experiment.

At the beginning the $BKI$ for each of the samples is calculated as the geometric average of the gene expression values in this particular sample. Since typically no previous knowledge is available about the analyzed genes at this stage of the experiment, it was assumed that the $BKI$ could be calculated based on all genes in the dataset as:

$$BKI[s] = \sqrt[N]{\prod_{n=1}^{N} D_{log}[n,s]} \tag{11}$$

Then, the Pearson correlation coefficient ($r$) with BKI is calculated for each gene:

$$r[n] = \sum_{s=1}^{S} \frac{(BKI[s] - \overline{BKI}) \cdot (D_{log}[n,s] - \overline{D_N}[n])}{SD_{BKI} \cdot SD_{DN}[n]} \tag{12}$$

where:

$$\overline{BKI} = \frac{\sum_{s=1}^{S} BKI[s]}{S} \tag{13}$$

$$\overline{D_N}[n] = \frac{\sum_{s=1}^{S} D_{log}[n,s]}{S} \tag{14}$$

$$SD_{BKI} = \sqrt{\frac{\sum_{s=1}^{S} (BKI[s] - \overline{BKI})^2}{S}} \tag{15}$$

$$SD_{DN}[n] = \sqrt{\frac{\sum_{s=1}^{S} (D_{log}[n,s] - \overline{D_N}[n])^2}{S}} \tag{16}$$

Finally, to maintain consistency with the other algorithms, the Stability Index value is calculated so that the lower it is, the more stable the considered gene:

$$SI[n] = 1 - r[n] \tag{17}$$

#### 1.2 NormFinder

The last normalization algorithm, which was examined, was called Normfinder. Its approach to finding the best reference genes is based on a mathematical model of gene expression, which, unlike other normalization algorithms, examines not only the overall suitability of candidate genes, but also studies the variations between groups in the dataset. Such groups can be found in a provided dataset if the biological samples represent cases of varying characteristics, eg. patients with a particular disease and healthy controls.

According to the model used as the foundation of this normalization algorithm, final stability measure value is calculated by combining both intra- and inter-group variances. All of the gene expression values have to be log-transformed beforehand.

At the beginning corrected expression values are determined:

$$DC_{log}[n,s] = D_{log}[n,s] - \overline{D_N}[n,g] - \overline{D_S}[s] - \overline{D[g]} \tag{18}$$

where $\overline{D_N}[n,g]$ is the average expression of gene $n$ in group $g$, $\overline{D_S}[s]$ is the average expression over all genes in sample $s$, $\overline{D[g]}$ is the average expression of all genes in all samples in group $g$, such that:

$$\overline{D_N}[n,g] = \frac{\sum\limits_{s \in g} D_{log}[n,s]}{S_g} \tag{19}$$

$$\overline{D_S}[s] = \frac{\sum\limits_{n=1}^{N} D_{log}[n]}{N} \tag{20}$$

$$\overline{D[g]} = \frac{\sum\limits_{s \in g} \sum\limits_{n=1}^{N} D_{log}[n,s]}{N \cdot S_g} \tag{21}$$

where $S_g$ is the number of samples in group $g$.

The variation of every gene expression within each of the groups (intra-group variation) is then estimated:

$$\sigma^2[n,g] = \frac{\sum\limits_{s \in g} (DC_{log}[n,s] - \overline{DC_N}[n,g])^2}{(S_g - 1) \cdot (1 - \frac{2}{N})} \tag{22}$$

where:

$$\overline{DC_N}[n,g] = \frac{\sum\limits_{s \in g} DC_{log}[n,s]}{S_g} \tag{23}$$

The formula for intra-group variation estimation was modified with respect to the one proposed in the original article, because for certain expression values provided in the dataset the resulting variance value would be negative. This fact would contradict the agreed upon definition and would prevent further calculations due to mathematical constraints.

The second part of NormFinder algorithm regards the calculation of differences in expression values between samples from different groups (inter-group variation).

$$d[n,g] = \overline{D_N}[n,g] - \widetilde{D_N}[n] - \widetilde{D_G}[g] + \widetilde{D} \tag{24}$$

where:

$$\widetilde{D_N}[n] = \frac{\sum\limits_{g=1}^{N_G} \overline{D_N}[n,g]}{N_G} \tag{25}$$

$$\widetilde{D_G}[g] = \frac{\sum\limits_{n=1}^{N} \overline{D_N}[n,g]}{N} \tag{26}$$

$$\widetilde{D} = \frac{\sum\limits_{g=1}^{N_G} \sum\limits_{n=1}^{N} \overline{D_N}[n,g]}{N \cdot N_G} \tag{27}$$

Ultimately, the final measure for gene expression stability takes into consideration both intra- and intergroup variability:

$$SI[n] = \rho[n] = \frac{\sum\limits_{g=1}^{N_G} \rho'[n,g]}{N_G} \tag{28}$$

with

$$\rho'[n,g] = \frac{\gamma \cdot |d[n,g]|}{\gamma + W[n,g]} + \sqrt{W[n,g] + \frac{\gamma \cdot W[n,g]}{\gamma + W[n,g]}} \tag{29}$$

where

$$W[n,g] = \frac{\sigma^2[n,g]}{S_g} \tag{30}$$

and

$$\gamma = max\left\{\frac{1}{(N-1)(N_G-1)}\sum_{n=1}^{N}\sum_{g=1}^{N_G} d[n,g] - \frac{1}{N \cdot N_G}\sum_{n=1}^{N}\sum_{g=1}^{N_G}\frac{\sigma^2[n,g]}{S_g}, 0\right\} \tag{31}$$

If the samples in the dataset are not divided into multiple groups, thus only one group is considered, there exists no inter-group variation and $d[n,g] = 0$.
